# Phylogenetic reclassification of vertebrate melatonin receptors to include Mel1d

**DOI:** 10.1101/574384

**Authors:** Elsa Denker, Lars O. E. Ebbesson, David G. Hazlerigg, Daniel J. Macqueen

## Abstract

The circadian and seasonal actions of melatonin are mediated by high affinity G-protein coupled receptors (melatonin receptors, MTRs), classified into phylogenetically distinct subtypes based on sequence divergence and pharmacological characteristics. Three vertebrate MTR subtypes are currently described: MT1 (MTNR1A), MT2 (MTNR1B), and Mel1c (MTNR1C / GPR50), which exhibit distinct affinities, tissue distributions and signaling properties. We present phylogenetic and comparative genomic analyses supporting a revised classification of the vertebrate MTR family. We demonstrate four ancestral vertebrate MTRs, including a novel molecule hereafter named Mel1d. We reconstructed the evolution of each vertebrate MTR, detailing genetic losses in addition to gains resulting from whole genome duplication events in teleost fishes. We show that Mel1d was lost separately in mammals and birds and has been previously mistaken for an MT1 paralogue. The genetic and functional diversity of vertebrate MTRs is more complex than appreciated, with implications for our understanding of melatonin actions in different taxa. The significance of our findings, including the existence of Mel1d, are discussed in an evolutionary and functional context accommodating a robust phylogenetic assignment of MTR gene family structure.

## INTRODUCTION

Melatonin is an ancient eukaryotic signalling molecule that regulates diverse biological functions. While best known as a regulator of biological rhythms in humans, this hormone also regulates energy balance, temperature, behavior, blood pressure, and seasonal reproduction. Melatonin is secreted by the pineal gland and targets the brain as well as peripheral tissues (Hardeland *et al.* 2011, Slominski *et al.* 2012), but is also produced by several tissues, eliciting paracrine effects (Weaver and Reppert 1990). The actions of melatonin depend on the spatiotemporal expression of high-affinity melatonin receptors (MTR), representing a specific class of G protein-coupled receptor (GPCR).

Three paralogous MTR family members have been characterized in jawed vertebrates, namely MT1 (Mel1a / MTNR1A), MT2 (Mel1b / MTNR1B), and Mel1c (MTNR1C / GPR50 in mammals) (Reppert *et al.* 1994, 1995a, 1995b). Despite showing overlap in expression, these different MTRs have evolved unique functions. MT1 has a higher affinity for melatonin than MT2 (Dubocovich and Markowska 2005), and in mammals, Mel1c has lost the ability to bind melatonin (Dufourny *et al.* 2008), though it does modulate melatonin signaling via its association with MT1 (Levoye *et al.* 2006). While MT1 associates with a range of G proteins to activate several distinct signalling pathways, eliciting wide-ranging cellular effects (Witt-Enderby *et al.* 2003), MT2 associates with a single G protein (Jockers *et al.* 2008). Owing to such functional divergence, different MTRs may have very distinct biological effects, even when expressed in the same cell types (*e.g.* Dubocovich and Markowska 2005).

A past study demonstrated melatonin binding in the brain of jawed vertebrates and lamprey, but not in hagfishes or amphioxus (Vernadakis *et al.* 1998). Thus, it is likely that high-affinity MTRs were present in the vertebrate ancestor, and were secondarily lost in some jawless fishes, as noted for several other traits (*e.g.* reduction of vertebrae-like elements - Ota et al. 2011; *Dlx* genes - Sugahara *et al.* 2013; reviewed in Kuraku 2013). MTR-like GPCR genes have also been discovered in urochordates, cephalochordates, hemichordates and echinoderms (Kamesh *et al.* 2008, Nordstrom *et al.* 2008, Krishnan *et al.* 2013), but their evolutionary affinity to the vertebrate MTRs remains ambiguous. The distinct MTRs of jawed vertebrates potentially originated during two rounds (2R) of whole genome duplication (WGD) at the stem of vertebrate evolution (*e.g.* Dehal and Boore, 2005), though this is yet to be established. Additional expansions in the MTR family of fishes (*e.g*. Shang & Zhdanova 2007; Hong *et al.* 2014) may owe to a further round of teleost-specific WGD (‘Ts3R’) in the common teleost ancestor, or additional lineage-specific WGD events in some lineages, *e.g.* the salmonid-specific 4R (‘Ss4R’) (Macqueen and Johnston, 2014; Lien *et al.* 2016), though, again, this has not been properly explored.

The overarching goal of this study was to re-examine the evolutionary history of vertebrate MTRs, using data in publically-available sequence databases for robust phylogenetic and comparative genomic reconstructions. Our findings concretely demonstrate a fourth ancestral MTR (‘Mel1d’), along with teleost-specific expansions in MTR diversity, likely owing to Ts3R and Ss4R. With a new evolutionary framework in place we reinterpret findings on vertebrate MTR sequence divergence and expression from past studies. Overall, this study highlights substantial unexplored diversity in MTR signalling within vertebrates, pointing to new lines of investigation.

## MATERIAL AND METHODS

### Sequence and phylogenetic analyses

Amino acid sequences encoded by MTR or FAT protocadherin family member genes were collected from representative jawed vertebrate species with high-quality genome assemblies. Details of these sequences are given in Table S1 (MTR) and Table S2 (FAT), which include database accession numbers and nomenclature matching the findings of our phylogenetic analyses. As a start point for the analysis, MTR/FAT proteins of human (*i.e.* MT1/MT2/Mel1c/GPR50 or FAT1/2/3) were used as queries in BLASTp (Altschul *et al.* 1997) searches to identify homologues within the NCBI database (https://www.ncbi.nlm.nih.gov/). We also used the Ensembl genome browser (https://www.ensembl.org/) to collect MTR family proteins from several species, using the EnsemblCompara method (Vilella *et al.* 2009).

The sequences were aligned using MAFFT v.7 (Katoh and Standley, 2013) with default settings and subjected to quality filtering using GBlocks with default settings (Talavera and Castresana, 2007). Final alignments of 300 (MTR) and 2,540 (FAT) amino acid positions (Additional Dataset 1) were used for tree building, done using BY and ML (MTR) or just ML (FAT) methods. ML trees were generated using IQ-TREE (Nguyen *et al.* 2015) via the IQ-TREE webserver (Trifinopoulos *et al.* 2016), employing the best-fitting amino acid substitution model selected with ModelFinder (Kalyaanamoorthy *et al.* 2017) under the Bayesian information criterion. The best fit models were JTT+F+I+ G4 for MTR and JTT+G4+I for FAT, where ‘JTT’ is the general matrix of Jones *et al.* 1992, ‘+I’ includes empirical estimation of the proportion of invariant sites, ‘+F’ includes empirical estimation of amino acid frequencies and ‘+G4’ denotes estimation of the gamma distribution parameter with 4 rate classes. The stability of branching in the ML trees was assessed using 1,000 ultrafast bootstrap replicates, (Hoang *et al.* 2018). The BY analysis (MTR dataset) was done in BEAST v1.8.3 (Drummond *et al*. 2012), employing an uncorrelated relaxed clock model (Drummond *et al.* 2005) and a Yule speciation prior (Gernhard, 2008), along with the best-fitting substitution model selected by IQ-TREE. A Markov chain Monte Carlo (MCMC) chain of 50 million generations was generated and sampled every 5,000 generations. Convergence of the MCMC chain was assessed using Tracer v1.7.1 http://beast.bio.ed.ac.uk/tracer). A maximum clade credibility tree was generated in TreeAnnotator (Drummond *et al.* 2012) after removal of the first 10% sampled trees.

### Comparative genomic and sequence analyses

Synteny analyses were performed using Ensembl genome browser annotations via the Genomicus platform (Nguyen *et al.* 2018). These analyses were supplemented with data from NCBI GenBank for species not available in Ensembl. Gene prediction and annotation for *Lethenteron camtschaticum* was performed using FGENESH (Soloyvev *et al.* 2006). Comparative analyses of MTR family amino acid sequences was done using the final alignment described above (note: the Gblocks filtering step served to remove flanking regions outside the transmembrane/loop regions, which were unaltered). The sequence similarity of the proposed vestigial MTR-like pseudogenes identified in our synteny analyses was established using BLASTx within the NCBI database.

### Data Availability

Supplemental material described in the paper is available at Figshare: XXX. Fig. S1. ML phylogenetic analysis of MTRs in vertebrates. This analysis was done using IQ-TREE with a high-confidence alignment of eighty MTRs (300 amino acid positions; Additional Dataset 1) and the best-fitting amino acid substitution model (JTT+F+I+G4). Numbers on branches are bootstrap support values. Other details as in the Fig. 1 legend (see main text) Table S1 provides details of all protein sequences used for phylogenetic analyses of the vertebrate MTR family. Table S2 provides details of all sequences used for phylogenetic analyses of the vertebrate FAT protocadherin family

**Fig. 1.**
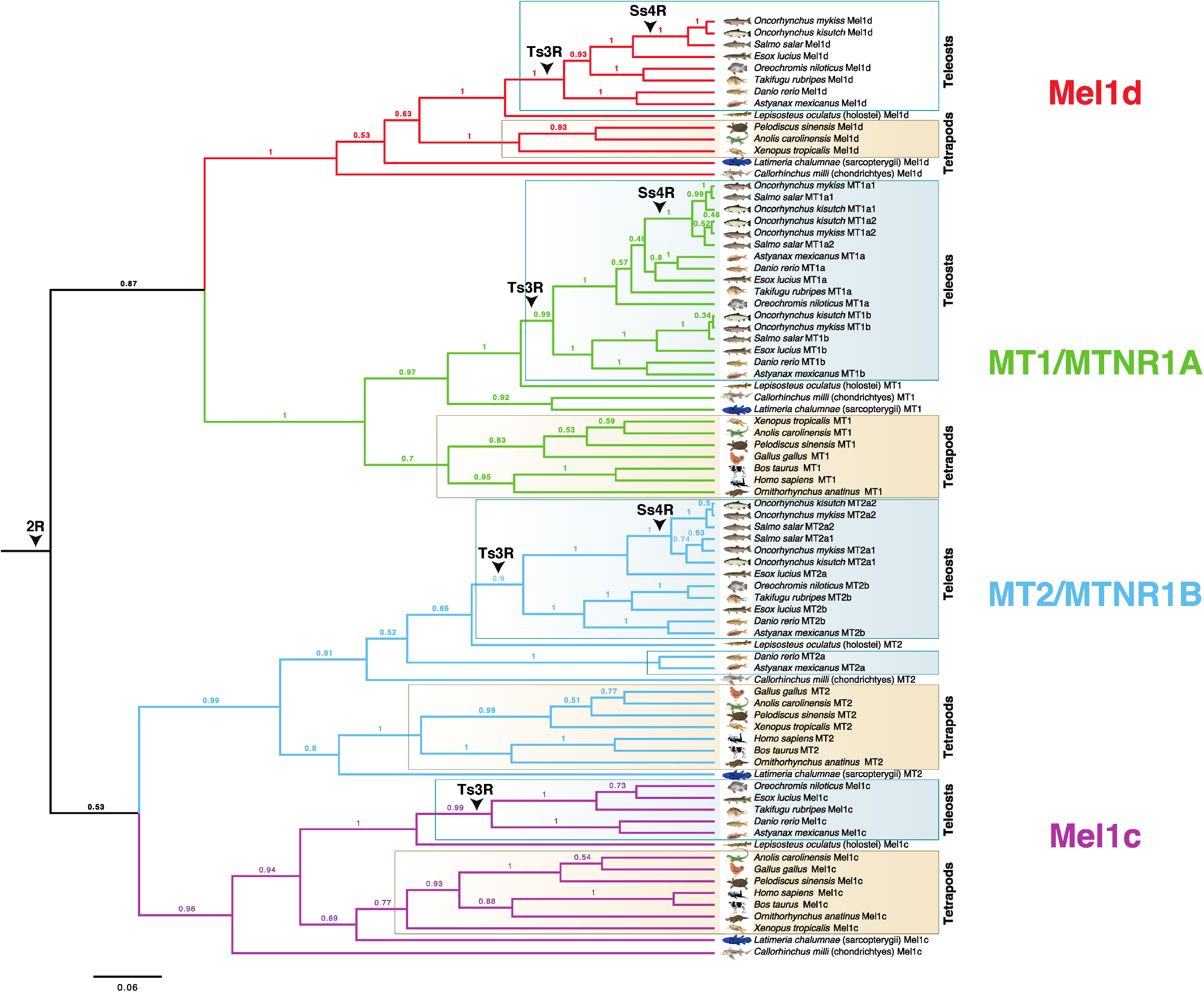
Bayesian phylogenetic tree of MTR family evolution in jawed vertebrates. The analysis was done using BEAST with a high-confidence alignment of eighty MTRs (300 amino acid positions; Additional Dataset 1), an uncorrelated relaxed molecular clock model and the best-fitting amino acid substitution model (JTT+F+I+G4). Numbers on branches are posterior probability support. Three WGD events in vertebrate evolution are shown (2R - ancestral to vertebrates; Ts3R - ancestral to teleosts; Ss4R - ancestral to salmonids). A ML tree was performed using the same data and is provided in Fig. S1.

Additional Dataset 1 is the MTR sequence alignment used for phylogenetic analysis and comparative sequence analysis. Additional Dataset 2 is the FAT alignment used for phylogenetic analysis.

## RESULTS

### Four MTRs are retained in jawed vertebrates

We identified eighty unique MTR family member proteins in sequence databases representing a standardized set of eighteen jawed vertebrate lineages (see MATERIALS AND METHODS). A Bayesian (BY) phylogenetic analysis was done (Fig. 1) incorporating a relaxed molecular clock model, which allows estimation of the most plausible root location in the tree (Drummond *et al.* 2006; *e.g.* Macqueen and Wilcox 2014; Redmond *et al.* 2018). Four distinct MTR clades (Fig. 1) had strong statistical support (posterior probability, PP: >0.96), and each was represented by cartilaginous fish, as well as ray-finned and lobe-finned fish lineages, with branching patterns closely matching expected species phylogeny (Fig. 1). Three of these clades correspond to known ancestral vertebrate MTR family members (*e.g.* Dufourny *et al.* 2008). The fourth clade is hereafter named ‘Mel1d’. The same four clades were strongly supported in an unrooted maximum likelihood (ML) phylogenetic analysis (bootstrap support: >96%) congruent with the BY tree (Fig. S1).

These analyses indicate that four distinct MTRs existed in the jawed vertebrate ancestor. However, the phylogenetic affinity of the four MTRs remains equivocal in the BY analysis, with moderate support for Mel1d/MT1 (PP: 0.87) and MT2/Mel1C (PP: 0.53) being paralogues, which can be explained parsimoniously by 2R (Fig. 1).

### Evolutionary history of individual vertebrate MTRs

Expanding on the above findings, we reconstructed a more detailed evolutionary history for each ancestral MTR in jawed vertebrates, accommodating gene losses, in addition to gains resulting from WGD events in teleosts (summarized in Fig. 2).

**Fig. 2.**
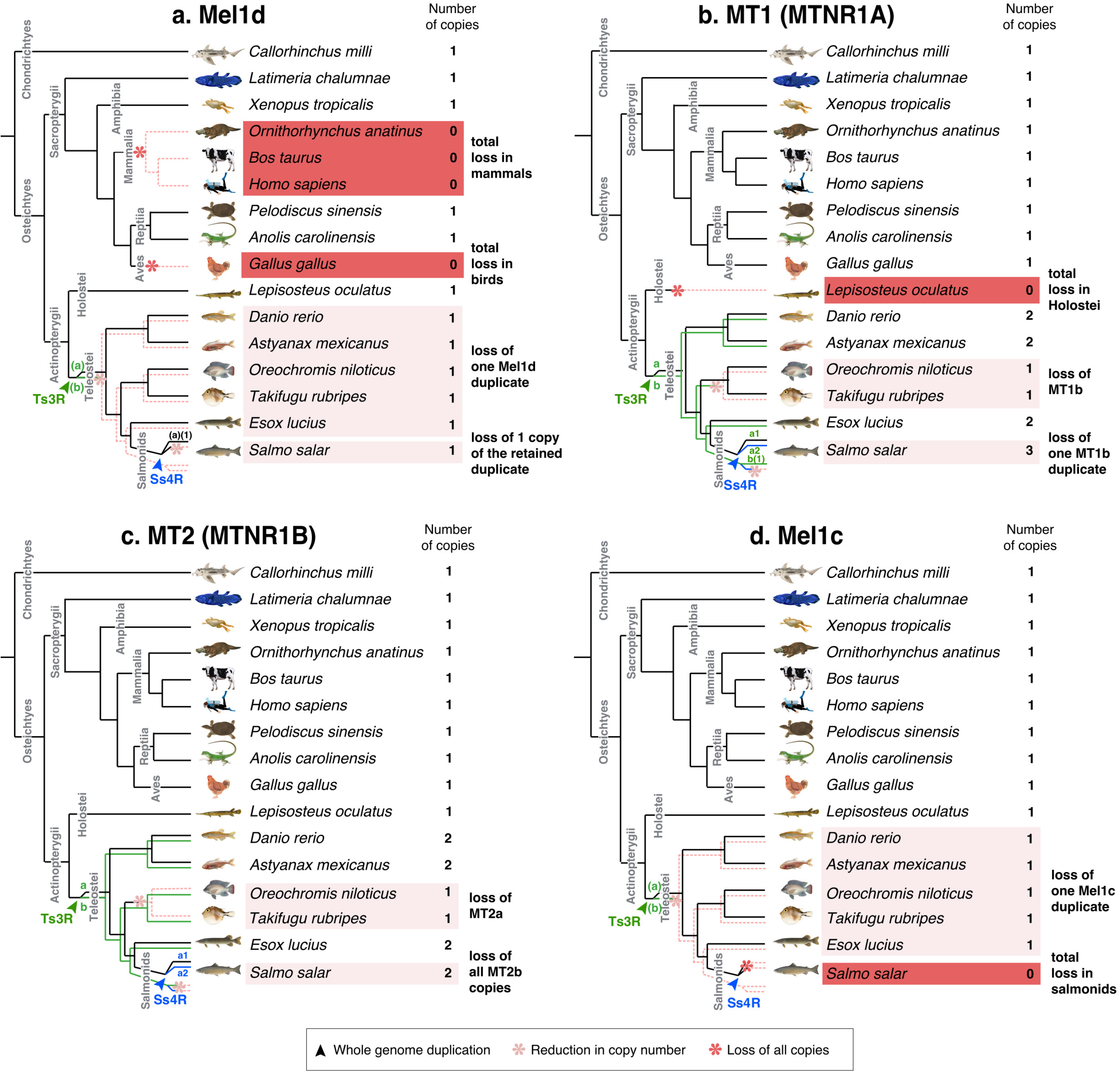
Proposed evolutionary history of each MTR family member, considering (a) Mel1d, (b) MT1, (c) MT2m, and (d) Mel1c. Species inferred to have lost all copies of a MTR gene are highlighted in dark red. Teleost species inferred to have lost paralogues of MTR genes arising from the Ts3R and Ss4R events are highlighted in light red.

Mel1d was encoded by a single gene in all represented species (Fig. 1, Fig. 2a) including teleosts, consistent with the loss of any paralogues created during Ts3R and Ss4R. In lobe-finned fish, Mel1d was identified in a coelacanth, an amphibian, and two reptiles, but was not identified in the mammals and birds represented in our trees (Fig. 2a). As only a small number of bird and mammals were included, we decided to search more broadly for Mel1d orthologues. Hence, BLAST searches of the complete set of proteins stored in NCBI for mammals (~4.6 million) and birds (~2.8 million) were done using reptile Mel1d orthologues as the query. Though hundreds of bird and mammal genomes are available in NCBI with protein-level annotations (spanning the diversity of each lineage), the top mammal/bird hit for reptile Mel1D was always MT1/MTNR1A (not shown). Considering our current understanding of amniote phylogeny (*e.g.* Chiari *et al.* 2012), our data requires that independent losses of Mel1d occurred in the ancestors to birds and mammals.

For all studied vertebrate species outside teleosts, we identified one copy of MT1, barring spotted gar, where MT1 was not identified (Fig. 1, Fig. 2b); its trace was retrieved in the genome after further analyses (see section below), representing a sequence annotated as a pseudogene. Several teleost species retain two or more ancestral MT1 copies (PP: 0.99, Fig. 1), which can be explained by Ts3R. These duplicates have been annotated in zebrafish as “Mtnr1aa” and “Mtnr1ab” (ZFIN 2008 - ZNC nomenclature, cloned as ‘“ZMel1a1” and “ZMel1a3” by Shang & Zhdanova 2007). Consequently, we maintained the same ‘a’ and ‘b’ nomenclature in all species according to inferences of orthology with zebrafish (Fig. 1). The two teleost-specific MT1 paralogues were not present in all teleost lineages, with MT1b absent from the studied acanthopterygians (tilapia and pufferfish). Salmonid-specific paralogues of MT1a (MT1a1 and 1a2) were identified, ancestral to three salmonid species (PP: 1.0, Fig. 2b), consistent with retention from Ss4R, though only a single copy of MT1b was retained in the same three species, suggesting ancestral loss following Ss4R (Fig. 1, Fig. 2b).

We identified one copy of MT2 in non-teleost vertebrate lineages, and evidence for teleost-specific paralogues (Fig. 2c). Two MT2 paralogues were identified in Ostariophysi members (zebrafish and Mexican cavefish) and northern pike (Protacanthoptergii); however, only one MT2 copy was identified in Acanthopterygii members (Nile tilapia and pufferfish) (Fig. 1, Fig. 2c). Branching patterns among these duplicates were not well resolved when considering species phylogeny. An ancestral teleost duplication event (*e.g*. Ts3R) predicts two paralogous MT2 teleost clades, each containing teleosts branching after expected species relationships (as seen for MT1a/b). However, a clade containing zebrafish “Mtnr1ba” (ZFIN 2008, “ZMel1b2” in Shang & Zhdanova 2007) branched outside other fish (including the non-teleost spotted gar) in both the BY and ML trees (Fig. 1 and S1). Internal to the spotted gar, there were two teleost MT2 clades, one containing zebrafish “Mtnr1bb” (ZFIN 2008, “ZMel1b1” in Shang & Zhdanova 2007) and other teleost lineages (northern pike and Acanthopterygii members), while the other contained a separate northern pike sequence and all MT2 sequences from salmonids. Given the strong support for the clade containing zebrafish “Mtnr1bb” (PP:1.0, Bootstrap support: 100%), we considered all sequences therein to be orthologous, and named them MT2b (to maintain the zebrafish “b” nomenclature) (Fig. 2c). We named the remaining teleost MT2 sequences as MT2a (Fig. 2c), under the hypothesis that orthology to zebrafish MT2a was obscured by a long-branch attraction artefact (note the long-branch length leading to Ostariophysi members for MT2a; Fig. S1). This scenario is parsimonious, as it allows for a single ancestral teleost duplication (*e.g*. Ts3R) rather than several lineage-specific MT2 gains. Accordingly, we propose that MT2a was lost in the ancestor to *Oreochromis* and *Takifugu*, while two salmonid duplicates of MT2a (MT2a1 and 2a2) were retained from Ss4R (Fig. 1 and S1, Fig. 2c). No copies of MT2b were identified in salmonids, suggesting a loss in the common salmonid ancestor (Fig. 2c).

As shown elsewhere (Dufourny *et al.* 2008), Mel1c and mammalian GPR50 proteins grouped together in a well-supported clade (Fig. 1). A single Mel1c copy was identified in all teleosts barring salmonids, which evidently lack Mel1c (Fig. 2d). This is consistent with a scenario where one Mel1c paralogue was lost early following Ts3R, and an additional loss occurred in the common salmonid ancestor (Fig. 2d).

### Synteny analysis supports phylogenetic assignment of vertebrate MTRs

Next, to gain an independent line of evidence to support our phylogenetic reconstructions, we compared the genomic regions harboring MTR-encoding genes among a range of vertebrate lineages. The local gene neighborhood containing each MTR family member was more or less conserved across jawed vertebrate evolution, defining identifiable synteny groups specific to each ancestral MTR (Fig. 3), including teleost and salmonid-specific paralogues (Fig. 4). The genomic neighborhood containing the single *MTR* locus of lampreys did not conserve synteny with an equivalent region containing any single MTR gene in gnathostomes. Instead, the genes surrounding the single *MTR* locus of lampreys showed notable similarity to a combination of genes located around the various gnathostome MTRs (Fig. 4e). This lends support to an ancestral origin of MTRs in the vertebrate lineage, but does not allow us to pinpoint the relationship of lamprey MTR to the four MTR family members of jawed vertebrates. One possible interpretation is that the duplications generating four gnathostome MTR genes occurred after the cyclostomes and gnathostomes split, with the lamprey genomic neighborhood reflecting a derived representation of the ancestral vertebrate state. However, the current consensus is that at least one round of WGD is shared by cyclostomes and gnathostomes (*e.g.* Kuraku *et al.* 2009, Stadler *et al.* 2004). In this scenario, conserved synteny between a single genomic region in the former to multiple blocks in the latter may be explained by one or more shared duplications followed by lineage-specific rediploidization, as proposed by Robertson *et al.* 2017.

**Fig. 3.**
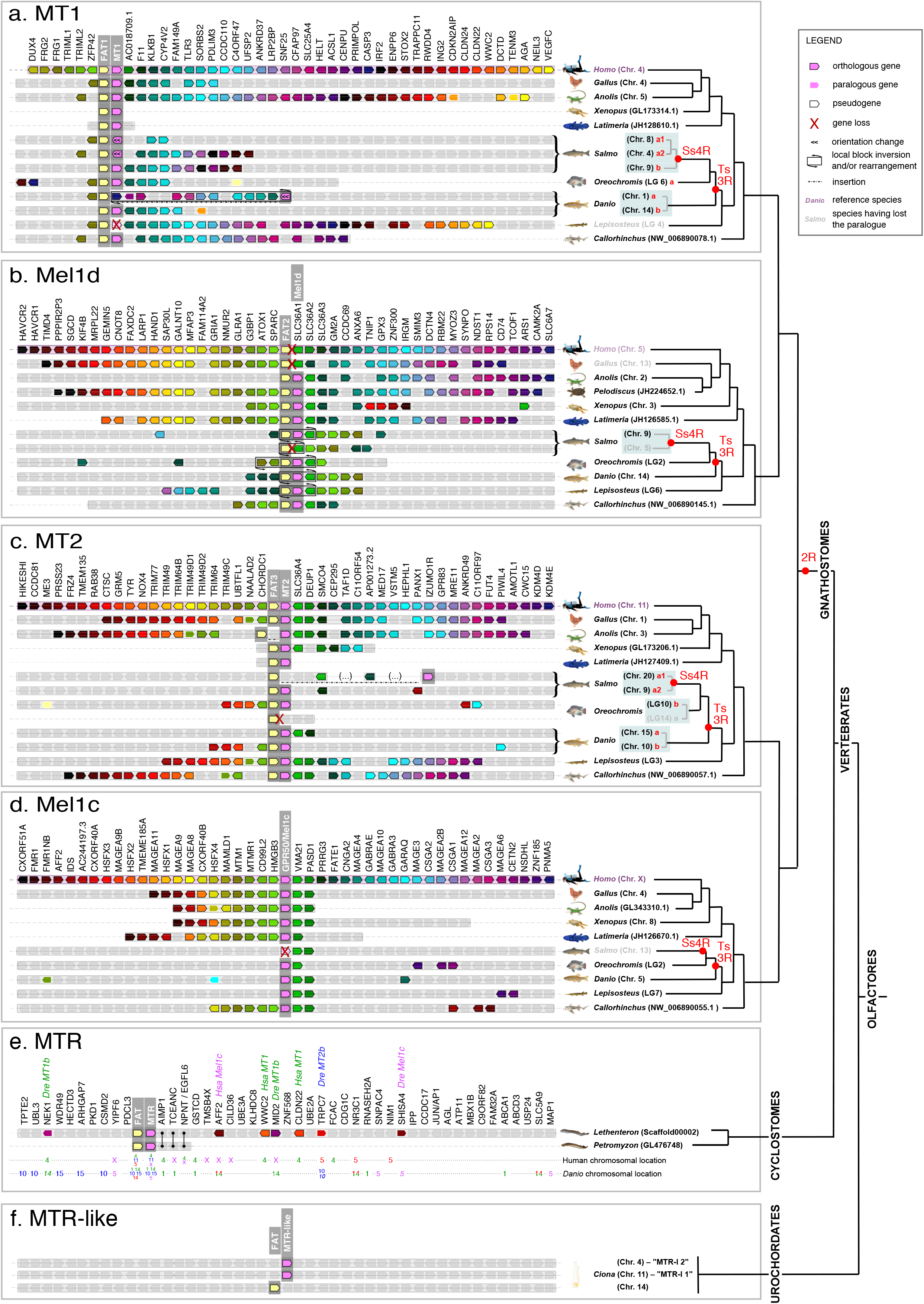
Conserved synteny between the genomic neighbourhood containing MTR orthologues of different lineages, shown for (a) jawed vertebrate MT1, (b) jawed vertebrate Mel1d, (c) jawed vertebrate MT2, (d) jawed vertebrate Mel1c, (e) comparing MTR from two lamprey species with jawed vertebrates, and (f) comparing a urochordate with vertebrates.

**Fig. 4.**
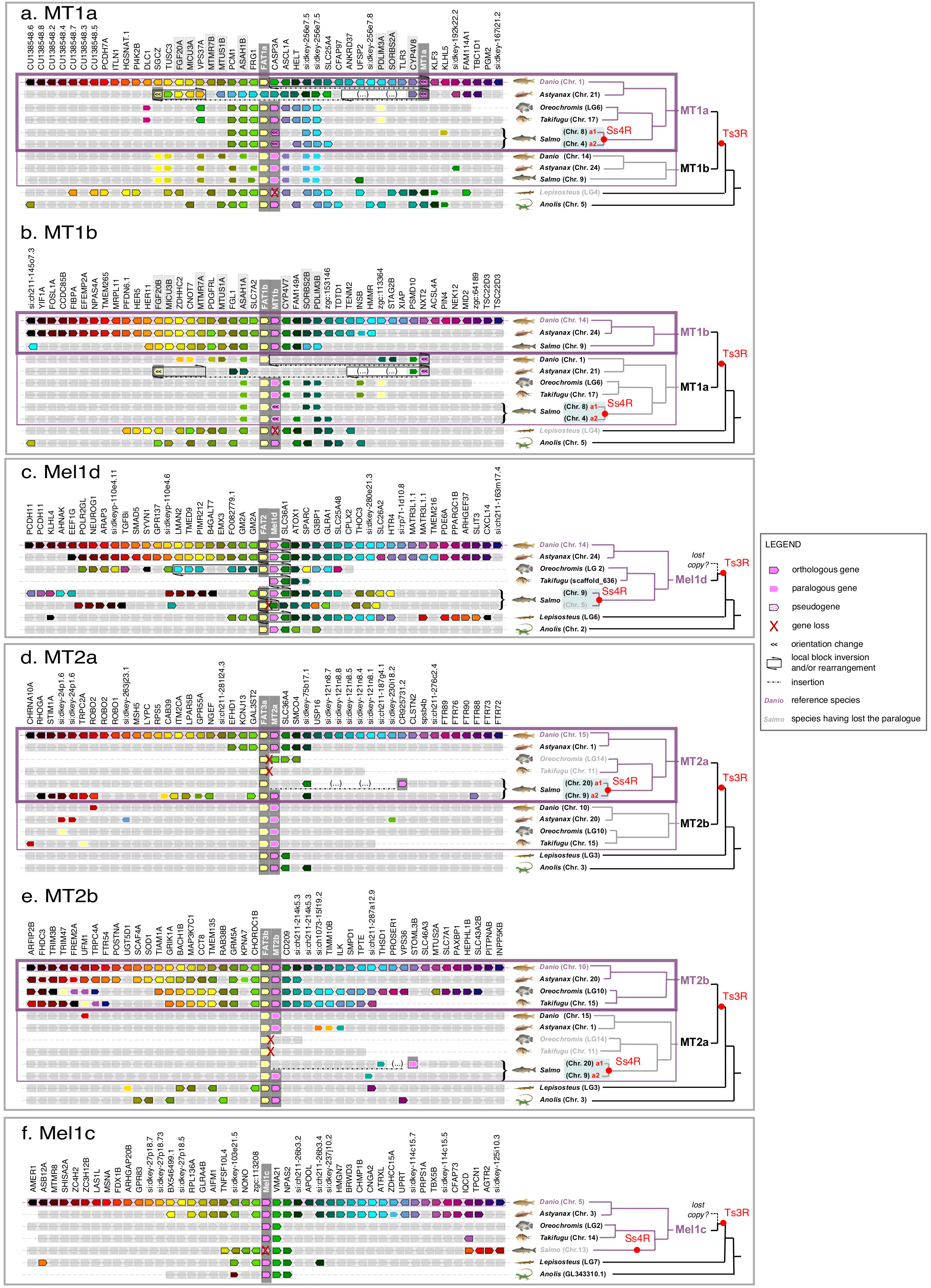
Conserved synteny between the genomic neighbourhood containing MTR paralogues retained from Ts3R and Ss4R, shown for (a) MT1a, (b) MT1b, (c) Mel1d, (d) MT2a, (e) MT2b, and (f) Mel1c.

### Genetic linkage between *MTR* and *FAT* genes

Tandem-linked MTR and *FAT* protocadherin gene family members are strongly conserved in all vertebrates (Fig. 3, Fig. 4). Specifically, *MT1*, *Mel1d*, and *MT2* were almost always in tandem with *FAT1*, *2*, and *3*, respectively (Fig. 3, Fig. 4). This association was absent for *Mel1c*, in addition to MTR co-orthologues from a sea squirt (Fig. 3f) and the Florida lancelet (not shown), defining this as a vertebrate-specific feature. Past studies have noted genetic linkage between MTR and FAT genes. For example, the FAT3-MT2 locus is involved in diabetes risk, with several SNPs involved in disease located between the two genes, implying potential functional links (*e.g.* Prokopenko *et al.* 2009, Dupuis *et al.* 2010). While, the reason for co-evolution of these loci is yet to be determined, the tandem organization of FAT and MTR genes indicates selective pressure to maintain an association that may be underpinned by a conserved feature of vertebrate physiology.

FAT family sequences also provide an independent source of phylogenetic information that may help reconstruct the evolution of the genomic regions containing linked MTR genes. In an ML analysis performed with FAT proteins from representative vertebrate species, we observed three clades (FAT1, 2 and 3) that branched according to expected species relationships (Fig. 5). When the ML tree was midpoint rooted, FAT1 (linked to MT1) and FAT2 (linked to Mel1d) were sister groups (Fig. 5), consistent with the sister relationship of MT1 and Mel1d recovered by the MTR phylogeny. Further, the teleost duplications observed for MTR genes were clearly identifiable in the respective tandem FAT genes (Fig. 5). Finally, the well-supported branching of salmonid FAT3a sequences with zebrafish FAT3a (i.e. linked to the *MT2a* gene, Fig 3c) adds weight to the hypothesis that salmonid/pike MT2 sequences are orthologous to zebrafish *MT2a* (Fig. 2c).

**Fig. 5.**
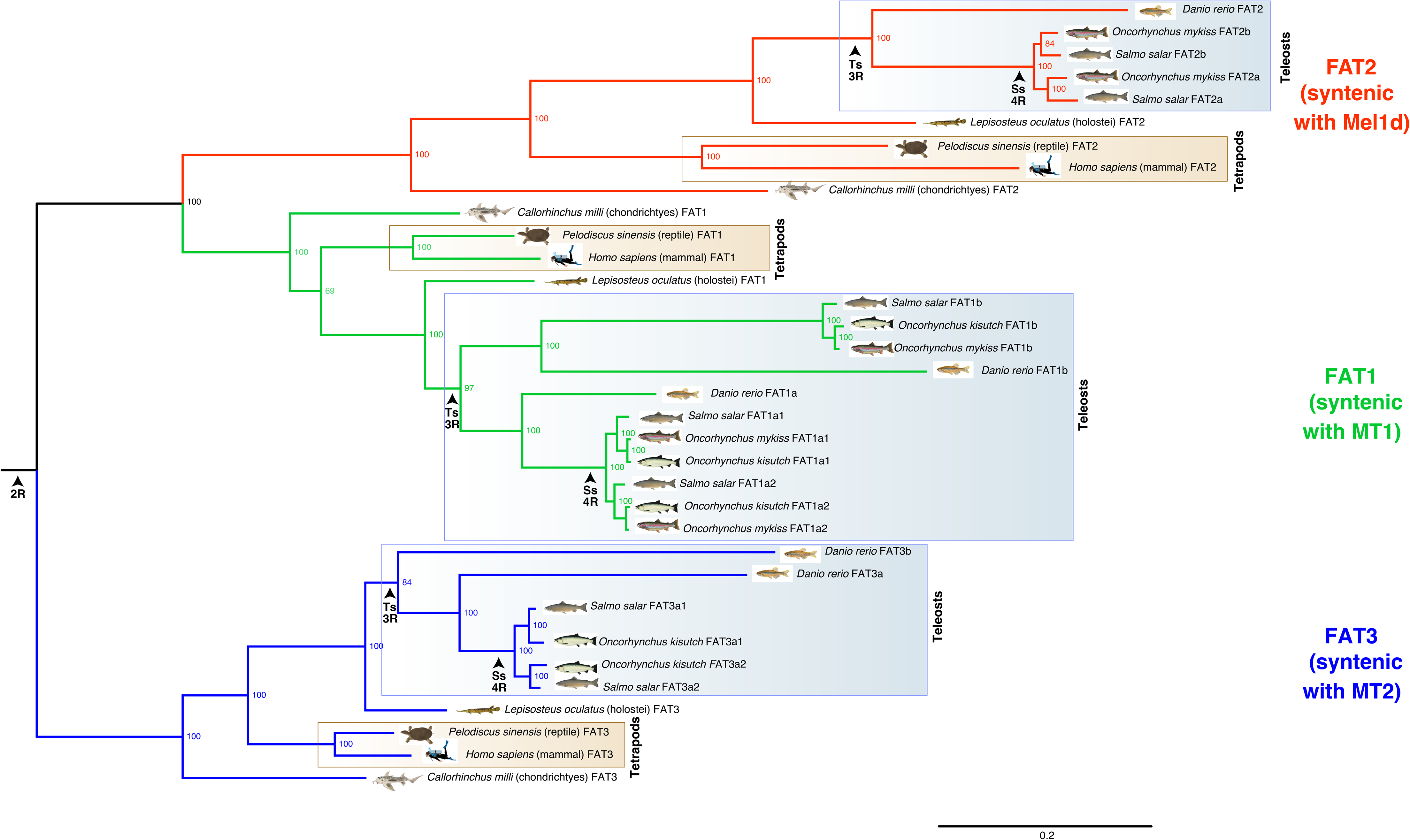
ML phylogenetic analysis of FAT atypical protocadherins in jawed vertebrates. The analysis was done using IQ-TREE with a high-confidence alignment of thirty-five FAT proteins (2,540 amino acid positions; Additional Dataset 2) and the best-fitting amino acid substitution model (JTT+G+I). Numbers on branches are bootstrap support values. Other details are as in the Fig. 1 legend.

### Synteny analyses support MTR losses

The conservation of synteny across vertebrate taxa in genomic regions containing MTR genes provides useful information on MTR genes inferred to be absent in sequence databases. In this respect, we observed that the genomic regions containing *Mel1d* in reptiles, frogs and fishes have matched syntenic regions in the human and chicken genomes (Fig. 3d). Consequently, the regions predicted to contain *Mel1d* in human and chicken have been properly assembled and are otherwise well annotated, consistent with *bone-fide* genetic losses of *Mel1d* in these species. The same approach allowed us to detect a pseudogene likely to be a vestige of *Mel1c* in Atlantic salmon (LOC106568030) (Fig. 3d), and a gene annotated as ‘non-coding’ bearing similarity with *MT1* (according to BLAST) at the predicted MT1 locus in spotted gar (LOC107077181) (Fig. 3a). Further, a second *FAT2* paralogue was detected in Atlantic salmon, supporting our previous conclusion of an ancestral loss of one Mel1d copy following Ss4R. Similarly, a second *FAT3* paralogue was detected in *Oreochromis*, non-paired with an *MTR2* gene (Fig. 3c), confirming the loss of *MT2a* in this species.

### Comparative sequence analysis of Mel1d with other MTRs

Having established that Mel1d is an ancestral vertebrate MTR, we sought to compare the primary amino acid sequence of this molecule to other MTR family members, hoping to gain clues on its function considering existing literature (Fig. 6).

**Fig. 6.**
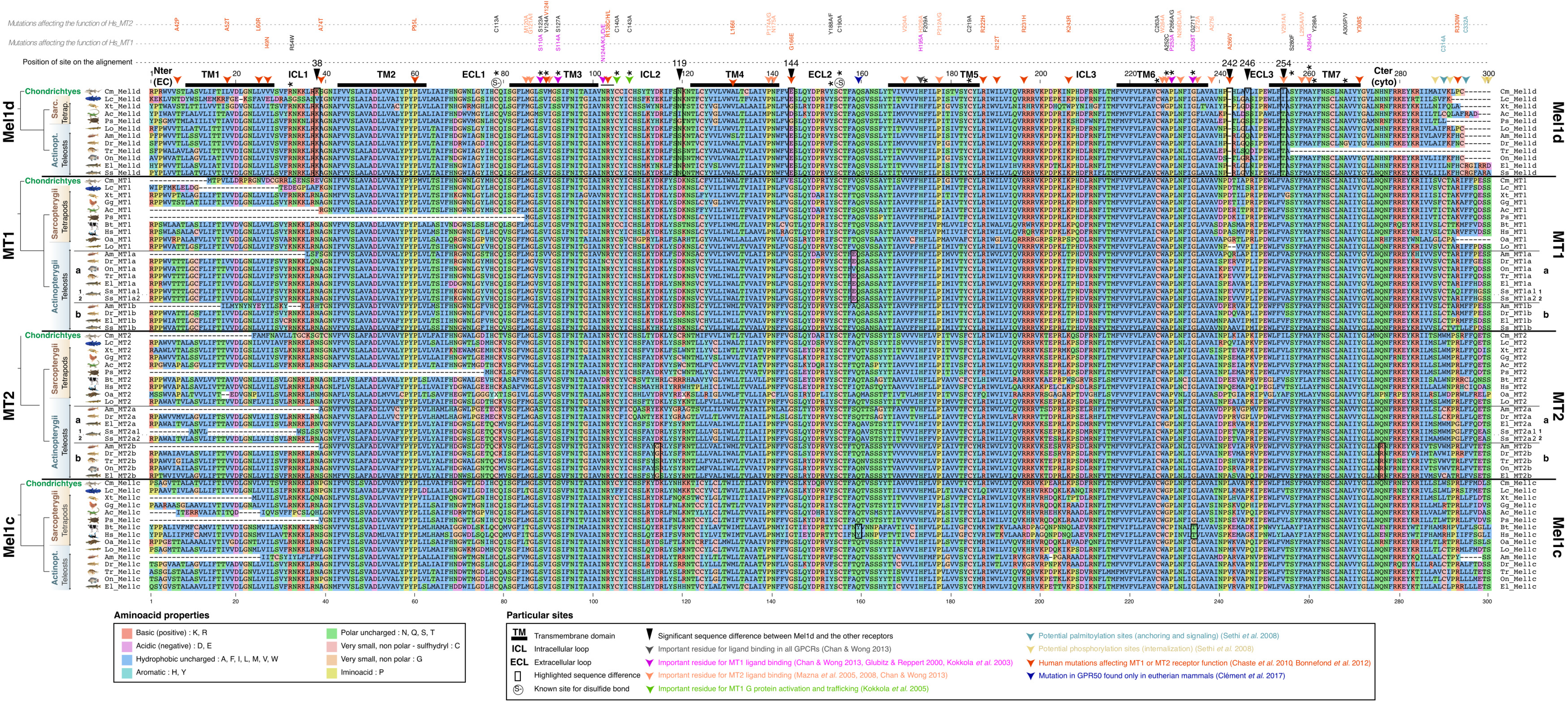
Alignment used to compare amino acid positions among vertebrate MTR proteins (matching to the alignment used for phylogenetic analysis; Additional Dataset S1). Species abbreviations: Ac = *Anolis carolensis* (green anole lizard); Am = *Astyanax mexicanus* (Mexican cavefish); Bt = *Bos taurus* (cattle); Cm = *Callorhinchus milli* (elephant shark); Dr = *Danio rerio* (zebrafish); El = *Esox lucius* (northern pike); Gg = *Gallus gallus* (chicken); Hs = *Homo sapiens* (human); Lc = *Latimeria chalumnae* (coelacanth); Lo = *Lepisosteus oculatus* (spotted gar); Oa = *Ornithorhynchus anatinus* (platypus); On = *Oreochromis niloticus* (Nile tilapia); Ps = *Pelodiscus sinensis* (Chinese softshell turtle); Tr = *Takifugu rubripes* (tiger pufferfish); Xt = *Xenopus tropicalis* (western clawed frog). Detailed annotation of sequences flagged up in the main text are provided within the figure.

We first examined the MTR transmembrane domains and ligand-binding residues, which have known functional importance. The characteristic seven transmembrane domain structure (TMDs) of all MTRs, critical for GPCR structure and ligand binding (Baldwin 1994), were conserved in Mel1d, MT1, MT2 and Mel1c (Fig. 6). Indeed, most of the residues identified as key for melatonin binding are readily identifiable in the Mel1d transmembrane domains (Fig. 6), in particular TM3, 6 and 7 (Gubitz & Reppert 2000, Kokkola *et al.* 2003, Mazna *et al.* 2005, 2008, Chan & Wong 2013). The only notable difference in the TMDs was that several Mel1d orthologues had threonine replacements at position 254, specific to this MTR. This position is important for melatonin binding in MT2 (valine-291 on human MT2), which was not reported for MT1 (Mazna *et al.* 2005). Outside the TMDs, two additional melatonin binding residues (asparagine-102 of the conserved NRY motif and alanine-238) were conserved in Mel1d (Fig. 6). Interestingly, a mutation in the second extracellular loop of GPR50 linked to the loss of melatonin binding function in mammals (Clément *et al.* 2017) was absent in Mel1d (Fig. 6).

Other key sites conserved in Mel1d included cysteine-78 and cysteine-155, responsible for a conserved disulfide bridge essential to MTR structure (Fig. 6). In addition, residues important for G protein activation and trafficking of MT1 (Kokkola *et al.* 2005) were all conserved in Mel1d (green arrows on Fig. 6). Putative palmitoylation site in MT1 and MT2 (cysteine-314 in MT1 and cysteine-332 in MT2, Sethi *et al.* 2008) required for G-protein interaction (light blue arrow on Fig. 6) were either not identified (MT2 cysteine-332) in Mel1d or absent from most species (MT1 cysteine-314). However a proximal conserved cysteine in position 294 of Mel1d (Fig.

6) may fulfil a similar function. Several phosphorylation sites have been suggested in the C-terminal cytoplasmic tail of MT1 and MT2, which might be important for β-arrestin-dependent receptor internalization (Ebisawa *et al.* 1994, Sethi *et al.* 2008, yellow arrows on Fig. 6). One of these sites is present on Mel1d, at position 288, however only in coelacanth and tetrapods. None of the other phosphorylation sites are present because of the shorter length of Mel1d, and this could be linked to differences in phosphorylation properties.

### Residue changes distinguishing Mel1d from other MTRs

The above analyses confirm that Mel1d has most of the canonical residues for melatonin binding and MTR structure/function. We next sought to identify conserved differences between Mel1d and the other MTRs, as candidates to impart functional properties unique to Mel1d.

Five extracellular or intracellular positions in Mel1d show substantial differences with either one or all other MTRs (Fig. 6). In most Mel1d orthologues, the first extracellular loop contains lysine (positive charged) at position 38, which is typically asparagine (neutrally charged) in the other MTRs. At position 144, which is almost always fixed as glycine in MT1, MT2 and Mel1c, Mel1d orthologues retain glutamic acid or aspartic acid. This replacement is presumed functionally significant, as glycine provides high conformational flexibility (Betts and Russell 2003), while glutamic and aspartic acid are highly negatively charged. At position 246, MTRs usually conserve proline (except for the two derived GPR50 from mammals), but Mel1d shows a high diversity of residues with diverse functional properties, suggesting a distinct mode of selective pressure. In the same loop (position 242), a gap is observed in all Mel1d sequences at an amino acid position that is variable among the other MTRs. Finally, a notable difference between Mel1d and MT1 is observed in position 119, in the second intracellular loop. Most MT1 sequences have aspartic acid at this position, while Mel1d conserves asparagine or serine, leading to a major difference in charge.

## DISCUSSION

Our unequivocal demonstration of a new ancestral vertebrate MTR forces a revision of current models for the origin and diversity of MTRs, and has biological implications for vertebrate lineages conserving distinct MTR gene repertoires.

It seems important to ask why Mel1d has previously been missed as a unique MTR, when the gene is readily detectable in sequence databases. This is likely partly due to a historic assumption that the MTR gene family structure of birds and mammals (i.e. MT1, MT2 and Mel1c) is representative for all vertebrates. Mel1d has high similarity with MT1, and has tended to be named ‘mtnr1a-like’ in genome databases. In addition, previous phylogenetic studies of MTRs have been based on small datasets (*e.g.* Reppert *et al.* 1995a; Mazurais *et al.* 1999; Park *et al.* 2006, 2007a,b; Shang & Zhdanova 2007; Hong *et al.* 2014), with biases in the taxa sampled, and could not by design distinguish Mel1d and MT1. A single past study noted a *Xenopus* MTR sequence that did not group with MT1, MT2 or Mel1c and concluded the existence of a novel MTR (Mel1d) (Shiu *et al.* 1996); correctly according to our findings. Our study benefits from a much broader survey of vertebrate MTR sequences, allowing us to conclude that Mel1d is at least 450 million years old, having been present in the jawed vertebrate ancestor.

Our phylogenetic reconstruction of MTRs will help the field going forwards, as researchers can be certain of which family member (including teleost-specific paralogues) they are studying, allowing more reliable conclusions in comparative studies of function and gene expression. We show that teleost-specific paralogues of MT1 are easily distinguished from Mel1d and provide a scheme to allow researchers to match teleost MTRs formerly named under several nomenclature systems to a single phylogenetically-assigned naming system accommodating orthologues and paralogues (Table 1).

**Table 1.** Phylogenetic assignment of teleost MTRs to a standardized nomenclature system.

| Species |  | Receptors (names attributed in the literature) and orthology group assignment from this study |  |  |  |  |  | References |
| --- | --- | --- | --- | --- | --- | --- | --- | --- |
|  |  | MT1 |  | MT2 |  | Mel1c | Mel1d |  |
|  |  | MT1a | MT1b | MT2a | MT2b |  |  |  |
| Dr | *Danio rerio*<br>(zebrafish) | **Z1.7**<br>(U31822.1) |  |  | **Z2.6**<br>(U31824.1) |  | **Z1.4**<br>(U31823.1) | Reppert *et al.* 1995(b) |
|  |  | **zMel1a1, Z1.7-4, mtnr1aa**<br>(NM_131393.1) | **zMel1a3**<br>(XM_688989.6) | **zMel1b2, Z6.2, Mel1b-19, mtnr1ba**<br>(NM_131395.1) | **zMel1b1, Z2.6-4, mtnr1bb**<br>(NM_131394.1) | **zMel1c, Z2.3, mtnr1c**<br>(NM_001161484.1) | **zMel1a2, Z1.4, mtnr1al**<br>(NM_001159909.1) | Shang & Zhdanova 2007 |
| Om | *Oncorhynchus mykiss*<br>(rainbow trout) | **R1.7**<br>(AF156262.1) = MT1a2 |  |  |  |  | **R1.4**<br>(AF178538.1) | Mazurais *et al.* 1999<br>**!!!!!! N.B. : the R2.6 gene (AF178929.1) is a chimera between Mel1d and MT2b2** |
| EI | *Esox lucius*<br>(northern pike) |  |  | **P2.6**<br>(AF188871.1) |  |  | **P1.4**<br>(XM_010903666.1) | Gaildrat and Falcón, 2000 |
| Oke | *Oncorhynchus keta*<br>(chum salmon) | **mel1a**<br>(AY356364.1) = MT1a2 |  |  |  |  | **mel1b**<br>(AY356365.1) | Shi *et al.* 2004 |
| Sg | *Siganus guttatus*<br>(golden rabbitfish) | **Mel1a**<br>(DQ768087.1) |  |  |  | **Mel1c**<br>(DQ768088.1) | **Mel1b**<br>(DQ522314.1) | Park *et al.*, 2006, 2007a,b, 2014 |
| Ca | *Carassius auratus*<br>(goldfish) | **Mel1a1.7**<br>(AB378058.1) |  |  | **Mel1b**<br>(AB378059.1) | **Mel1c**<br>(AB378060.1) | **Mel1a1.4**<br>(AB378057.1) | Ikegami *et al.* 2009 |
|  |  | **G1.7**<br>(AB481372.1) |  | **G6.2**<br>(AB481374.1) | **G2.6 type1**<br>(AB481373.1) | **Mel1c**<br>(AB481375.1) | **G1.4**<br>(AB481371.1) | Saito, unpublished |
| DI | *Dicentrarchus labrax*<br>(sea bass) | **dIMT1**<br>(EU378918.1) |  |  | **dIMT2**<br>(EU378919.1) | **dIMel1c**<br>(EU378920.1) |  | Sauzet *et al.*, 2008<br>Herrera-Pérez P *et al.*, 2010 |
| Sse | *Solea senegalensis*<br>(Senegal sole) |  |  |  | **ssMT2**<br>(FM213464.1) | **ssMel1c**<br>(FM213465.1) | **ssMT1**<br>(FM213463.1) | Confente *et al.* 2010 |
| On | *Oreochromis niloticus*<br>(Nile tilapia) | **mel1a**<br>(AY569971.1) |  |  |  |  |  | Jin *et al.*, 2013 |
| Ec | *Epinephelus coioides*<br>(orange-spotted grouper) | **MT1**<br>(JX524508.1) |  |  | **MT2**<br>(JX524509.1) |  |  | Chai *et al.*, 2013 |
| Bp | *Boleophthalmus pectinirostris*<br>(mudskipper) | **Mtnr1a1.7**<br>(KC622030.1) |  |  | **Mtnr1b**<br>(KC622031.1) | **Mtnr1c**<br>(KC622032.1) | **Mtnr1a1.4**<br>(KC622029.1) | Hong *et al.* 2014 |
| Tn | *Takifugu niphobles*<br>(grass puffer) | **mel1a1.7**<br>(AB492764.1) |  |  | **mel1b**<br>(AB492765.1) | **mel1c**<br>(AB492766.1) | **mel1a1.4**<br>(AB492763.1) | Ikegami *et al.* 2015 |
| Pn | *Porichthys notatus*<br>(plainfin midshipman - "singing" fish) | **mtnr1A1.7**<br>(HQ007044) |  | **Mel1b**<br>(KT878765.1) |  |  | **mtnr1a1.4**<br>(HQ007045) | Feng & Bass, 2016, Feng unpublished |
| Amel | *Amphiprion melanopus*<br>(cinnamon clownfish) | **MT-R1**<br>(HM107821.1) |  |  |  |  |  | Choi *et al.*, 2016 |

*Notes: Phylogenetic assignment according to findings of this study; previous publications using distinct nomenclature systems are provided. Sequences in red signal a significant change in assignment.*

### Insights into Mel1d function: reinterpreting expression data in teleosts

While not being previously recognized as a unique vertebrate MTR, Mel1d has already been studied in various teleosts (Table 1). These past studies demonstrate that the *Mel1d* transcript is abundantly expressed in a manner like other MTR family members, but showing differences that may underlie unique functions. A pattern seems conserved across multiple species, where Mel1d and MT1a expression is higher in brain and retina, respectively (*e.g.* Park *et al.* 2006, 2007a,b; Ikegami *et al.* 2009: Confente *et al.* 2010; Hong *et al.* 2014). Mel1d tends to be more strongly expressed in brain regions associated with visual perception (*e.g.* Mazurais *et al.* 1999; Gaildrat and Falcón, 2000; Shi *et al.* 2004; Confente *et al.* 2010; Hong *et al.* 2014). Many peripheral tissues were reported to express Mel1d with species-specific differences and in a distinct manner to other MTRs (Park *et al.* 2006, 2007a,b; Ikegami *et al.* 2009; Confente *et al.* 2010; Hong *et al.* 2014). Such data suggests involvement of Mel1d in photoreceptive processes, along with broader regulatory roles in the physiological functions of peripheral organs.

Rhythmical oscillations in the expression of Mel1d have also been reported, with variations depending on species, organ and season. In zebrafish, a day/night oscillation of MTR brain gene expression (peaking at night) was noted for all six MTR paralogues, including Mel1d, with further expression upregulation in response to melatonin administration (Shang & Zhdanova 2007). In golden rabbitfish, MT1a, Mel1d and Mel1c expression was higher at night for brain and retina, with Mel1d levels peaking at different times (Park *et al.* 2006, 2007a,b, 2014). In goldfish, Mel1d was the only MTR showing rhythmical oscillations in optic tectum expression, while the same was true for MT1a in retina, both peaking at the night-day transition (Ikegami *et al.* 2008). In a marine pufferfish, Mel1d, MT1a and Mel1c showed synchronous daily cycling of expression in the pineal gland with a nocturnal peak (Ikegami *et al.* 2015). Conversely, in golden rabbitfish pineal gland, oscillations were desynchronized for the same three MTRs (Park *et al.* 2006, 2007a,b). Daily rhythmicity in Mel1d expression has also been observed in peripheral tissues (liver and kidney) of golden rabbitfish, with higher expression during the day, opposite to the brain/retina (Park *et al.* 2006, 2007b). In addition to daily variation in regulation, Mel1d expression is regulated by other cycles, for example showing semilunar oscillation in the diencephalon of mudskipper (Hong *et al.* 2014) and ultradiurnal oscillation in a marine pufferfish, which may be circatidal (Ikegami *et al.* 2015). Mel1d expression in the Senegalese sole exhibited stronger day-to-night and seasonal variation than other MTR family members, with reciprocal differences recorded between retina and optic tectum (Confente *et al.* 2010). Therefore, past work shows that Mel1d is regulated during multiple biological cycles in teleosts, showing variations distinct from other MTRs, implying functional distinctiveness.

### Functional divergence between Mel1d and MT1?

High protein-level similarity between Mel1d and MT1, taken with the conservation of all key residues in the MTR transmembrane domains, strongly implies that Mel1d binds melatonin. Notably, residues showing conserved replacements between Mel1d and MT1 are all located in extracellular or cytoplasmic loops, which is predicted to impact interactions with other proteins, in particular signalling partners, rather than melatonin. Strikingly, one of these sites corresponds to a documented human MT1 mutation studied *in vitro* (Chaste *et al.* 2010). The replacement of glycine-144 (MT1) with glutamic acid or aspartic acid corresponds to a G166E mutation in human MT1, associated with impaired activation of cAMP signalling, despite retention of strong melatonin binding (Chaste *et al.* 2010). The elephant shark retains glutamic acid at this position in both MT1 and Mel1d, suggesting this represents the ancestral state, with functional divergence arising in the common ancestor to lobe and ray-finned fishes. It is also intriguing to observe that Mel1d of two tetrapods have apparently reverted to glycine in this position, indicating selection towards the ancestral residue.

### Why was Mel1d lost in mammals and birds?

Further work will be needed to establish the extent of conservation in Mel1d function and regulation across different vertebrate lineages. This should focus on reptiles and amphibians, where the function of this gene has not been studied experimentally. Such studies may help explain the specific biological requirements for Mel1d, and reveal why the gene was lost independently in mammals and birds. It is notable that mammals and birds stand out from other vertebrates when considering their melatonin-dependent light detection and clock systems. Mammals have lost extraocular light perception and relocated control of their biological clock away from melatonin-producing pinealocytes to the suprachiasmatic nucleus (Falcón *et al.* 2009). Birds have both the ancestral pineal clock and melatonin production system, but also independently developed a clock system in the homologue of the suprachiasmatic nucleus and use retinal detection (Cassone 1991, Falcón *et al.* 2009). Another distinguishing feature specific to both groups is homeothermy, with modulatory effects of melatonin on body temperature regulation reported in humans (Cagnacci *et al.* 1992; Viswanathan *et al.* 1990) and Japanese quail (Underwood and Edmonds 1995). Extrinsic temperature variation appears a less important zeitgeber for the circadian clock of homeotherms relative to poikilotherms (Rensing and Ruoff, 2002), which are known to use melatonin to regulate behavioral thermoregulation (Lutterschmidt *et al.* 2003). In addition, birds and mammals are the only vertebrates that have evolved (through convergent mechanisms) stereotypical slow wave and rapid eye movement sleep phases, linked to melatonin regulation in mammals (Lesku *et al.* 2011). Such changes in the physiological role of melatonin and consequent re-organization of melatonin response pathways, may have been the ultimate driver for Mel1d redundancy and gene loss through relaxation of purifying selection.

Another melatonin-associated function that is present in vertebrate lineages retaining Mel1d (in addition to lamprey), but lost in both mammals and birds, is the negative regulation of pigmentation development in the dark, known as the “body-blanching response” (Hamasaki and Eder 1977, Norris and Carr 2013). In fishes, melatonin is thought to regulate chromatosome aggregation in different kinds of chromatophores (Fujii 2000); Mel1d is expressed in the skin of mudskipper (together with MT1 - Hong *et al.* 2014), the goldfish (together with MT2 and Mel1c - Ikegami *et al.* 2008) and the sole (together with MT2 - Confente *et al.* 2010). In addition, in sole skin, Mel1d is the only MTR to be up-regulated at night. It is therefore possible that Mel1d is involved in skin physiology and pigment regulation in fish chromatophores.

### Expansion of the MTR repertoire of teleosts

Contrary to mammals/birds, there has been a trend towards evolutionary expansion in the MTR repertoire of teleosts, as observed in many gene families with paralogues retained from Ts3R (Glasauer and Neuhauss, 2014) and Ss4R (Houston and Macqueen, 2019). Interestingly, not all MTR family members were affected equally. While we identified multiple paralogous copies of MT1 and MT2 - presumed to have been retained from Ts3R and Ss4R - Mel1c and Mel1d were always single copy, requiring repeated losses of paralogues generated during gene duplication or WGD events. This is compatible with a hypothesis where the functions or expression-level regulation of MT1 and MT2 can be divided among paralogous copies, following the well-established subfunctionalization model, or potentially reflects fixation of new adaptive functions among MT1/MT2 paralogues (Stoltzfus 1999 and Force *et al.* 1999). In this respect, we observed several amino acid substitutions between MT1a vs. MT1b and MT2a vs. MT2b (Fig. 6), consistent with protein-level functional divergence. Conversely, selection has operated in a distinct manner for Mel1c and Mel1d, with any duplicates generated being quickly purged by selection for reasons that remain to be established, but potentially linked to dosage constraints, or a mechanism of regulation that cannot be divided across distinct loci.

### Conclusions

Mel1d is one of four ancestral vertebrate MTRs that shows a wide phylogenetic distribution and has both conserved and divergent functional characteristics compared to MT1, MT2 and Mel1c, including at the protein-sequence level and in terms of expression linked to chronobiological traits. Additional work is needed to characterize the functional distinctiveness of Mel1d compared to other MTRs and to explain why unique MTR repertoires have been conserved in different vertebrate lineages.

## Conflict of interest

The authors declare no conflict of interest.

## Author contributions

The study was initiated by DH, LOEE and DJM. Sequence collection, alignment and phylogenetic analyses was done by DJM. Comparative genomic and sequence analyses were done by ED. The manuscript was written by ED and DJM, with contributions from DH and LOEE.

## Funding information

This work was funded by the Frimedbio (Fri prosjektstøtte for medisin, helse og biologi) program of the Research Council of Norway (grant number 241016 - “Light & Salt - Thyroid hormone deiodinase paralogues & the evolution of complex life-history strategy in salmonids”). DJM was supported from BBSRC Institute Strategic Programme funding to The Roslin Institute (grant ref: BBS/E/D/10002071).

